# Spatial cell firing during virtual navigation of open arenas by head-restrained mice

**DOI:** 10.1101/246744

**Authors:** Guifen Chen, John A King, Yi Lu, Francesca Cacucci, Neil Burgess

**Author notes:** equal contribution (Guifen Chen, John A King, Yi Lu).

## Abstract

We present a mouse virtual reality (VR) system which restrains head-movements to horizontal rotations, potentially compatible with multi-photon imaging. We show that this system allows expression of the spatial navigational behaviour and neuronal firing patterns characteristic of real open arenas (R). Place and grid, but not head-direction, cell firing had broader spatial tuning in VR than R. Theta frequency increased less with running speed in VR than in R, while firing rates increased similarly in both. Place, but not grid, cell firing was more directional in VR than R. These results suggest that the scale of grid and place cell firing patterns, and the frequency of theta, reflect translational motion inferred from both virtual (visual and proprioceptive) cues and uncontrolled static (vestibular translation and extra-maze) cues, while firing rates predominantly reflect visual and proprioceptive motion. They also suggest that omni-directional place cell firing in R reflects local-cues unavailable in VR.

## Introduction

Virtual reality (VR) offers a powerful tool for investigating spatial cognition, allowing experimental control and environmental manipulations that are impossible in the real world. For example, uncontrolled real-world cues cannot contribute to determining location within the virtual environment, while the relative influences of motoric movement signals and visual environmental signals can be assessed by decoupling one from the other^1,2^. In addition, the ability to study (virtual) spatial navigation in head-fixed mice allows the use of intracellular recording and two photon microscopy^3-12^. However, the utility of these approaches depends on the extent to which the neural processes in question can be instantiated within the virtual reality (for a recent example of this debate see^13^).

The modulation of firing of place cells or grid cells along a single dimension, such as distance travelled along a specific trajectory or path, can be observed as virtual environments are explored by head-fixed mice^2-4,6-9,12^ or body-fixed rats^14-16^. However, the two-dimensional firing patterns of place, grid and head-direction cells in real world open arenas are not replicated in these systems, in which the animal cannot physically rotate through 360^°^.

By contrast, the two-dimensional spatial firing patterns of place, head direction, grid and border cells have been observed in VR systems in which rats can physically rotate through 360^° 17,18^. Minor differences with free exploration remain, e.g. the frequency of the movement-related theta rhythm is reduced^17^, perhaps due to the absence of translational vestibular acceleration signals^14,30^. However, the coding of 2-d space by neuronal firing can clearly be studied. These VR systems constrain a rat to run on top of an air-suspended Styrofoam ball, wearing a “jacket” attached to a jointed arm on a pivot. This allows the rat to run in any direction, its head is free to look around while its body is maintained over the centre of the ball.

However, these 2-d VR systems retain a disadvantage of the real-world freely moving paradigm in that the head movement precludes use with multi-photon microscopy or intracellular recording. In addition, some training is required for rodents to tolerate wearing a jacket. Here we present a VR system for mice in which a chronically implanted head-plate enables use of a holder that constrains head movements to rotations in the horizontal plane while the animal runs on a Styrofoam ball. Screens and projectors project a virtual environment in all horizontal directions around the mouse, and onto the floor below it, from a viewpoint that moves with the rotation of the ball, following^17,18^. See Figure 1 and Materials and Methods.

**Figure 1.**
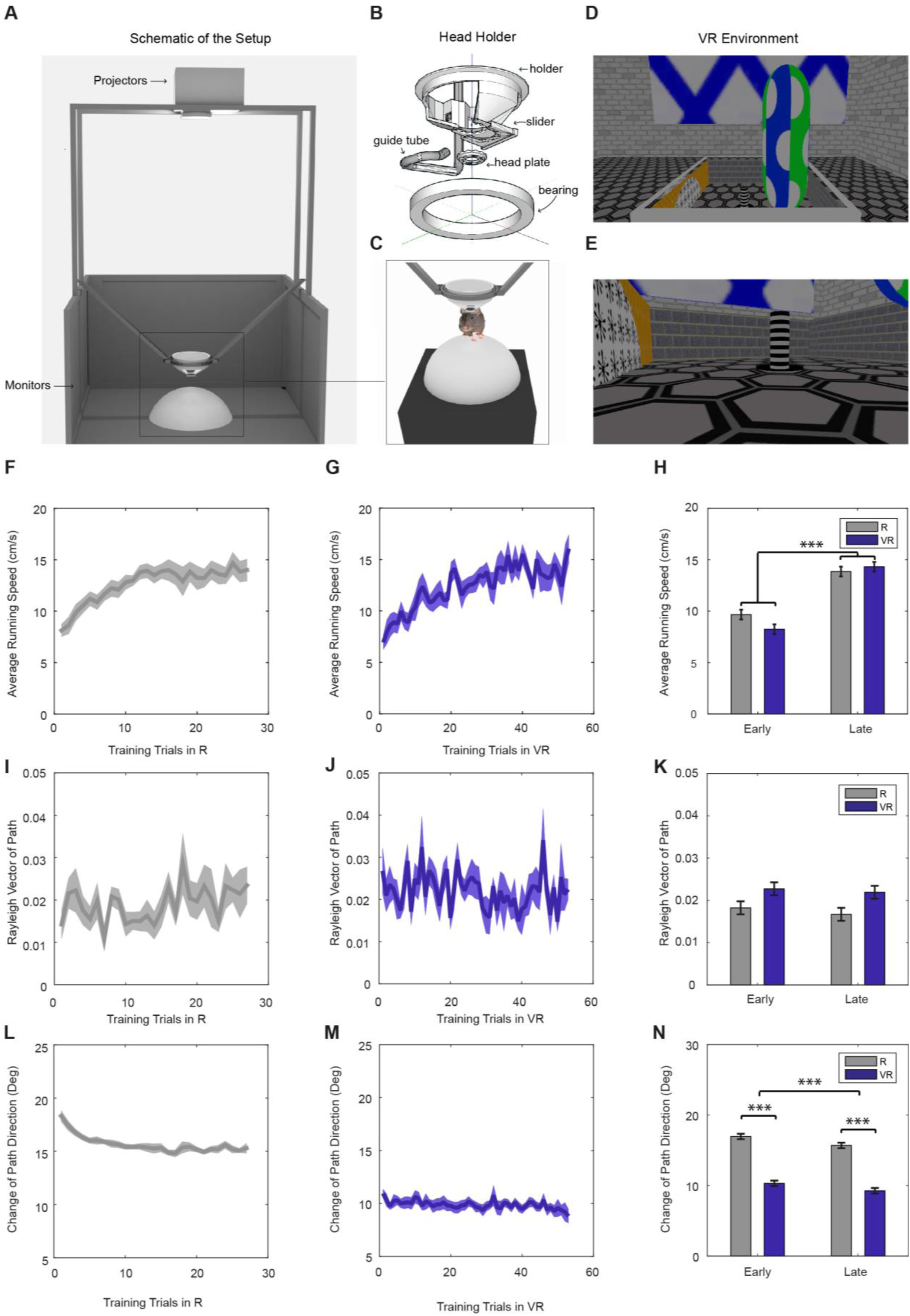
Virtual reality setup and behaviour within it. (A) Schematic of the VR setup. (B) A rotating head-holder. (C) A mouse attached to the head-holder. (D-E) Side views of the VR environment. (F-G) Average running speeds of all trained mice (n=11) across training trials in R (F) and VR (G) environments. (H) Comparisons of the average running speeds between the first 5 trials and the last 5 trials in both VR and R environments, showing a significant increase in both (n=11, p<0.001, F(1,10)=40.11). (I-J) Average Rayleigh vectors of running direction across training trials in R (I) and VR (J). (K) Comparisons of the average Rayleigh vectors of running direction between the first 5 trials and the last 5 trials in both VR and R. Directionality was marginally higher in VR than in R (n=11, p=0.053, F(1,10)=4.82) and did not change significantly with experience. (L-M) Average changes of running direction (absolute difference in direction between position samples) across training trials in R (L) and VR (M). (N) Comparisons of the changes of running direction between the first 5 and last 5 trials in both R and VR. Animals took straighter paths in VR than R (n=11, p<0.001, F(1,10)=300.93), and paths became straighter with experience (n=11, p<0.001, F(1,10)=26.82). Positions were sampled at 2.5Hz with 400ms boxcar smoothing in (I-N).

We demonstrate that this system allows navigation to an unmarked location within an open arena, showing that mice can perceive and remember locations defined by the virtual space. We also show that the system allows expression of the characteristic 2-dimensional firing patterns of place cells, head-direction cells and grid cells in electrophysiological recordings, making their underlying mechanisms accessible to investigation by manipulations of the VR.

## Results

### Navigation in VR

Eleven mice were trained in the virtual reality system (see Figure 1 and Materials and Methods). All training trials in the VR and the real square environments from the 11 mice were included in the behavioural analyses below. The mice displayed an initially lower running speed when first experiencing the real-world recording environment (a 60x60cm square), but reached a higher average speed after 20 or so training trials. The increase in running speed with experience was similar in the virtual environments (Figure 1F-H). Running speeds did not differ between the 60cm and 90cm virtual environments used for recording in 7 and 4 of the mice respectively (12.01±2.77 in 60cm VR, 14.33±4.19cm/s in 90cm VR, p = 0.29). Running directions in the VR environment showed a marginally greater unimodal bias compared to the real environment (R; Figure 1K). Mice displayed a greater tendency to run parallel to the four walls in VR, a tendency which reduced with experience (Figure S2). They also took straighter, less tortuous, paths in VR than in R, as would be expected from their head-fixation (Figure 1L-N).

In the fading beacon task, performance steadily improved across 2-3 weeks of training (Figure 2D, one trial per day). They learned to approach the fixed reward location and could do so even after it became completely unmarked (fully faded, see Figure 2 and the Supplementary video of a mouse performing the task, and Materials and Methods for details of the training regime).

**Figure 2.**
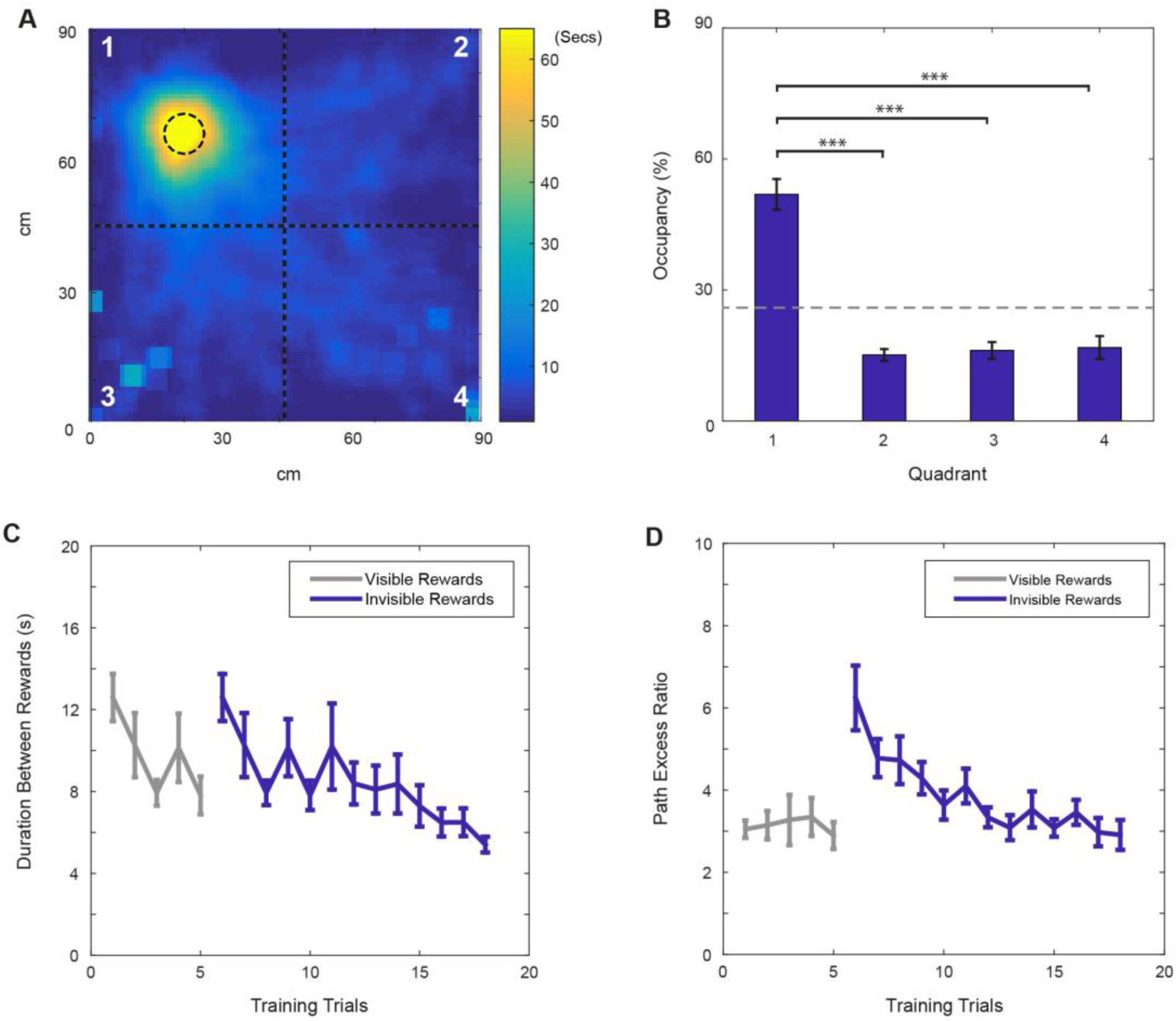
Performance on the ‘fading beacon’ task. (A) An example heat map showing the distribution of locations during a 40-min trial (mouse#987, trial#24). The dotted circle in the 1^st^ quadrant shows the location of the faded reward. (B) Average time spent (as % of total time) in each quadrant of the square (numbered in A) showed a clear bias (n=11, p < 0.001, F(3,30) = 39.03), with time spent in the 1^st^ quadrant was significantly higher than in the others (*** denotes significance at p<0.001, ** at p<0.01). (C) Average durations between the 3^rd^ and the 4^th^ rewards across training trials. (D) Average path excess ratios between the 3^rd^ and the 4^th^ rewards across training trials (means ± s.e.m). Note that in each set of four rewards, the 1^st^, 2^nd^ and 3^rd^ rewards appeared at random locations in the virtual square, marked by visual beacons, the 4^th^ reward was located at a fixed location. Grey lines show trials when the fixed-location rewards were marked by visual beacons. Blue lines show trials when the fixed rewards were not marked. See Supplementary video.

### Electrophysiology

We recorded a total of 231 CA1 cells from 7 mice: 175 cells were classified as place cells in the real environment, 186 cells in the virtual environment, and 154 cells were classified as place cells in both real and virtual environments (see Materials and Methods).

We recorded 141 cells in dorsomedial Entorhinal Cortex (dmEC) from 8 mice, 82 of them were classified as grid cells in the real environment, 65 of them were grid cells in the virtual environment, and 61 were classified as grid cells in both real and virtual environments. Among these 141 recorded cells, 16 cells were quantified as head-direction cells (HDCs) in R, 20 cells were HDCs in VR, with 12 cells classified as HDCs in both R and VR environments.

Place cells recorded from CA1 showed spatially localised firing in the virtual environment, with similar firing rates in the virtual and real square environments. Place cells had larger firing fields in VR than in R, by a factor 1.44 (field size in VR/ field size in R), which did not differ between those recorded in a 60×60cm versus a 90×90cm VR environment (1.44 in 90cm, 1.43 in 60cm, p=0.66). The spatial information content of firing fields in VR was lower than in R. In addition, the firing of place cells was more strongly directionally modulated in VR than in R. See Figure 3.

**Figure 3.**
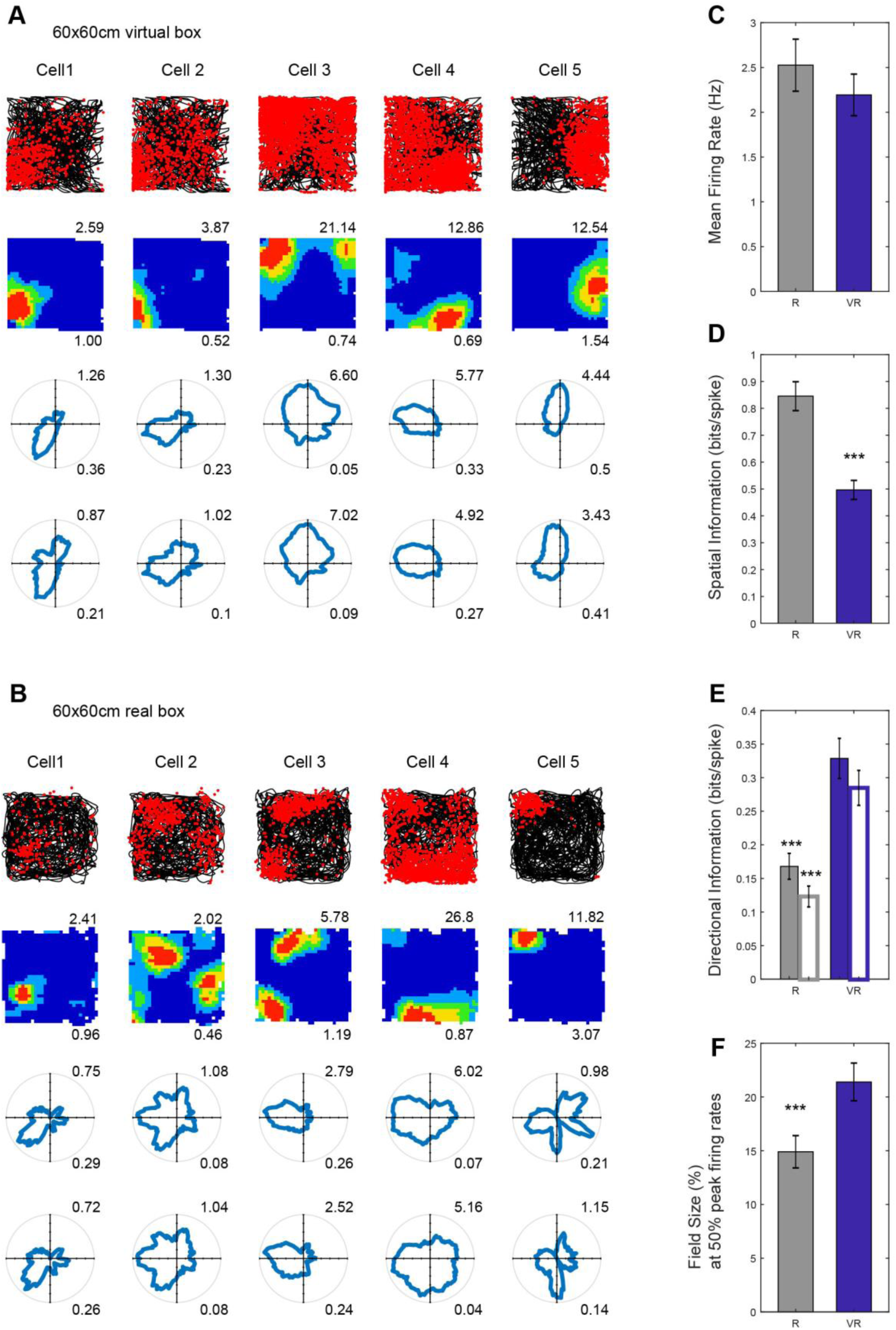
Place cell firing in real and virtual environments. (A-B) Five place cells simultaneously-recorded in a 60x60cm virtual square (A) and in a 60×60cm real square (B, one cell per column). Top row: 40-min running trajectory (black line) with red dots showing the locations of spikes; 2nd row, firing rate maps, maximum firing rate (Hz) shown at top right, spatial information rate(bits/spike) bottom right; 3^rd^ and 4^th^ row: polar plots of directional firing rates (3^rd^ row: standard binning; 4^th^ row: after fitting a joint ‘pxd’ model to account for inhomogeneous sampling), maximum firing rate shown at top right, directional information bottom right. (C) Comparison of mean firing rates. There was no significant difference between R and VR (n=154, t(153)=1.67, p=0.10). (D) Comparison of spatial information rates. Rates were significantly higher in R than in VR (n=154, t(153)=8.90, p<0.001). (E) Comparison of directional information rates using standard (solid bars) and pxd binning (open bars). Place cells were more directional in VR than in R (standard n=154, t(153)=6.45, p<0.001; pxd, n=154, t(153) = 7.61, p<0.001). (F) Comparison of field sizes between R and VR. The place fields included location bins with spikes higher than 50% of peak firing rates. Field sizes were calculated based on their proportion to the size of the test squares. The field sizes were bigger in VR than in R (n=154, t(153)=4.38, p<0.001). Blue bars indicate results from VR trials, grey bars indicate R trials.

One possible contribution to apparent directionality in firing could be inhomogeneous sampling of direction within the (locational) firing field. This can be controlled for by explicitly estimating the parameters of a joint place and direction (‘pxd’) model from the firing rate distribution^24^. However, using this procedure did not ameliorate the directionality in firing (see Figure 3). Further analyses showed that firing directionality increased near to the boundaries in both virtual and real environments (where sampling of direction is particularly inhomogeneous), but that the additional directionality in VR compared to R was apparent also away from the boundaries. See Figure S3.

Grid cells recorded in dmEC, showed similar grid-like firing patterns in VR as in R, with similar firing rates and ‘gridness’ scores. The spatial scale of the grids was larger in VR than in R, with an average increase of 1.42 (grid scale in VR/ grid scale in R, n=6 mice), which did not differ between those recorded in a 60x60cm versus a 90x90cm VR environment (1.43 in 60cm VR, 1.36 in 90cm VR, p=0.78). The spatial information content of grid cell firing was lower in VR than R, as with the place cells. Unlike the place cells, the grid cells showed only a slight increase in directionality from R to VR, which, unlike for place cells, appears to reflect inhomogeneous sampling of directions within firing fields, as the effect was not seen when controlling for this in a joint ‘pxd’ model. See Figure 4. It is possible that low directional modulation of the firing of a grid cell could reflect directionally modulated firing fields with different directional tuning. Accordingly we checked the directional information in the firing of each field, without finding any difference between R and VR (Figure 4H).

**Figure 4.**
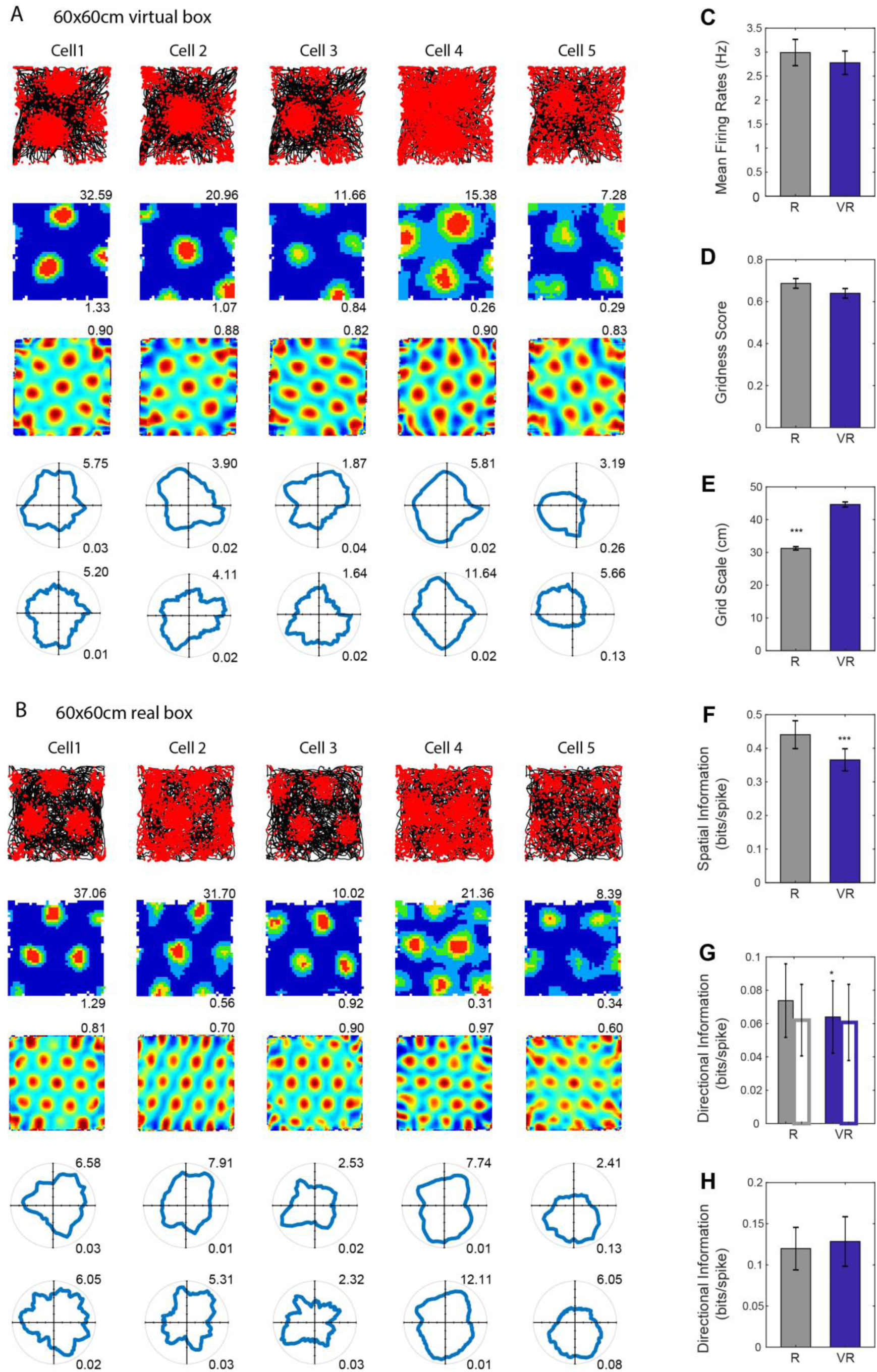
Grid cell firing in real and virtual environments. (A-B) Five grid cells simultaneously-recorded in a 60×60cm virtual square (A) and in a 60×60cm real square (B, one cell per column). Top row: 40-min running trajectory (black line) with red dots showing the locations of spikes; 2nd row, firing rate maps, maximum firing rate (Hz) shown at top right, spatial information (bits/spike) bottom right; 3^rd^ row: spatial autocorrelation maps, numbers on the top right show gridness scores; 4^th^ and 5^th^ rows: polar plots of directional firing rates (4^th^ row: standard binning; 5^th^ row: after fitting a joint ‘pxd’ model to account for inhomogeneous sampling), maximum firing rate shown at top right, directional information bottom right. (C) Comparison of mean firing rates. There was no significant difference between R and VR (n=61, t(60)=1.71, p=0.09). (D) Comparison of gridness scores, higher in R than VR but not significantly so (n=61, t(60)=1.67, p=0.10). (E) Comparison of grid scales, showing significantly larger scales in VR than in R (n=61, t(60)=15.52, p<0.001). (F) Comparison of spatial information in bits/spike, which was significantly higher in R than VR (n=61, t(60)=4.12, p<0.001). (G) Comparison of directional information. Grid fields tended to be slightly more directional in VR than in R (n=61, t(60)=2.04, p<0.05), although the bias disappeared when the rates were calculated based on pxd plots (open bars, n=61, t(60)=0.32, p=0.75). (H) Comparison of directional information in individual firing fields. There was no significant difference between the R and VR trials based on pxd plots (n=61, t(60)=0.53, p=0.60). Blue bars indicate results from VR trials, grey bars indicate R trials.

We also recorded head-direction cells in the dmEC, as previously reported in rats^21^ and mice^25^. These cells showed similar firing rates in VR and R, with similar tuning widths. See Figure 5. The relative differences in the tuning directions of simultaneously recorded head-direction cells was maintained between R and VR, even though the absolute tuning direction was not (see Figure S4).

**Figure 5.**
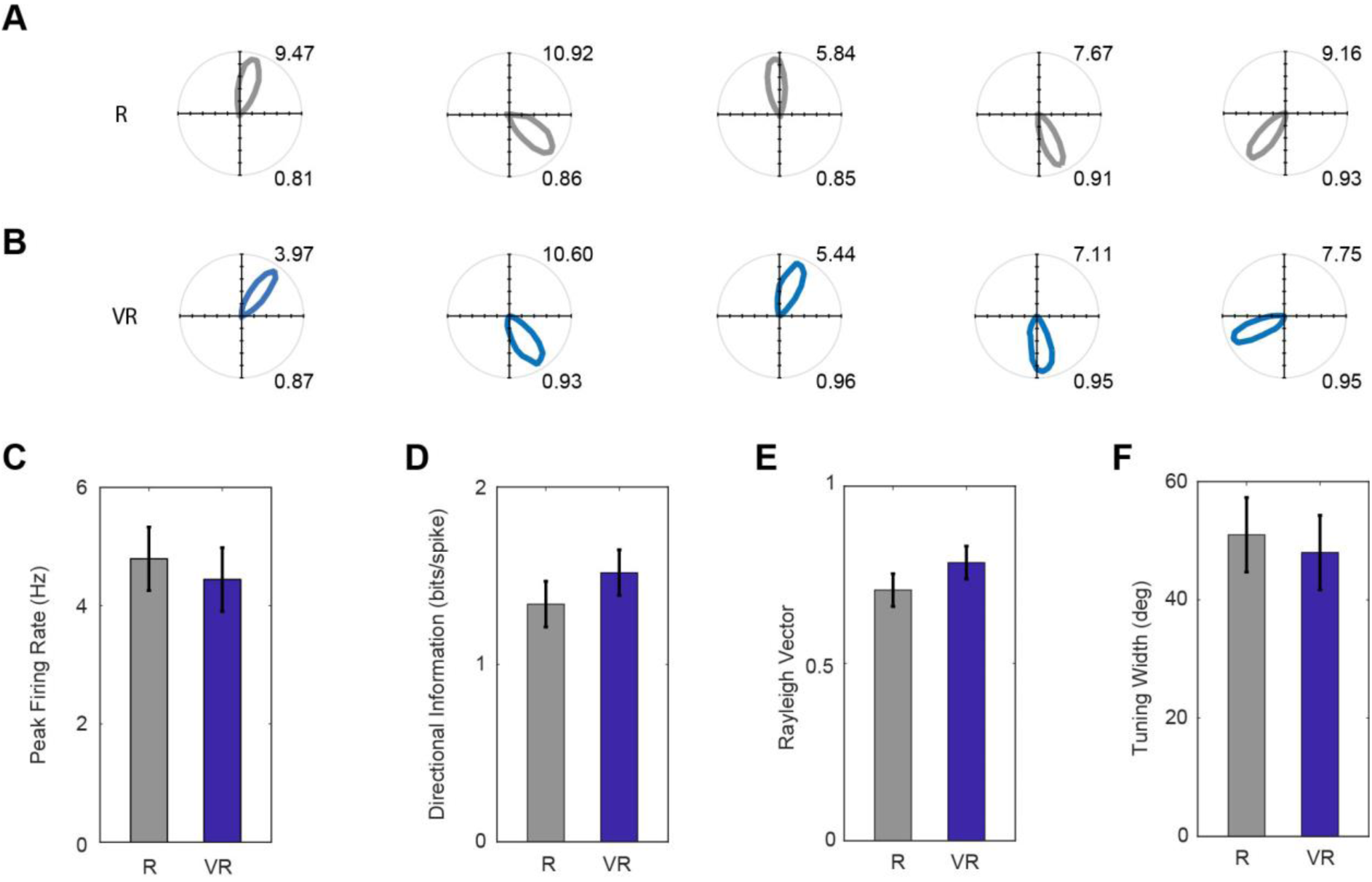
Head direction cell firing in real and virtual environments. (A-B) Polar plots of five simultaneously recorded HD cells in dmEC in VR (A) and R (B, one cell per column). Maximum firing rates are shown top right, Rayleigh vector length bottom right. (C-F) Comparisons of basic properties of HD cells in dmEC between R and VR. There were no significant differences in peak firing rates (t(11) = 0.65, p = 0.53; C); spatial information rate (t(11)=1.38, p=0.19; D); Rayleigh vector length (t(11)=1.69, p=0.12; E); and tuning width (t(11)=0.48, p=0.64; F).

The translational movement defining location within the virtual environment purely reflects feedback (visual, motoric and proprioceptive) from the virtual reality system, as location within the real world does not change. However, the animal’s sense of orientation might reflect both virtual and real-world inputs, as the animal rotates in both the real and virtual world. To check for the primacy of the controlled virtual inputs versus potentially uncontrolled real-world inputs (e.g. auditory or olfactory), we performed a 180^°^ rotation of the virtual environment between trials. Note that the geometry of the apparatus itself (square configuration of screens, overhead projectors on either side) would conflict with rotations other than 180^°^. In separate trials, we observed a corresponding rotation of the virtual firing patterns of place, grid and head-direction cells, indicating the primacy of the virtual environment over non-controlled real world cues. See Figure 6.

**Figure 6.**
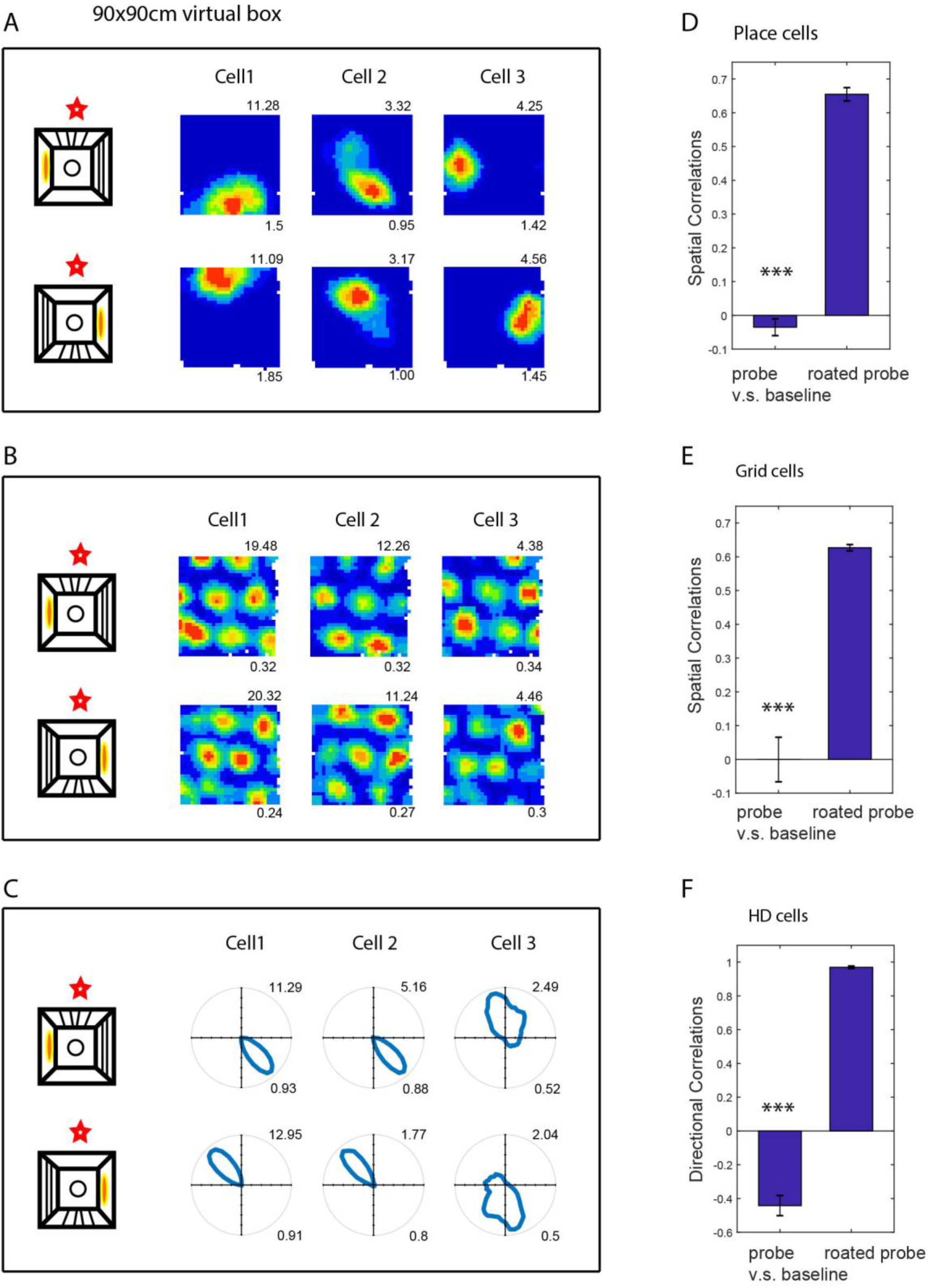
Effect of rotating the virtual environment on spatial firing patterns. (A-C) Three simultaneously recorded CA1 place cells (A), dmEC grid cells (B) and dmEC HD cells (C). The upper rows are the firing patterns in the baseline trials, and the lower rows are the patterns in the probe trials. The far-left column shows the schematic of the manipulation: all virtual cues were rotated 180^°^ relative to the real environment (marked by the red stars). Maximum firing rates are shown top right, spatial information (A), gridness (B) or Rayleigh vector length (C) bottom right. (D-F) Spatial correlations between the 180^°^ rotated probe trials and the baseline trials were significantly higher than between the probe trials and the baseline trials (spatial correlations for place cells, n=123, t(122)=19.44, p<0.001; grid cells, n=18, t(17)=9.41, p<0.001; HD cells, n=17, t(16)=24.77, p<0.001).

The animal’s running speed is known to correlate with the firing rates of cells, including place cells, grid cells and (by definition) speed cells^21,22,26^, and with the frequency of the local field potential theta rhythm^27-29^. So these experimental measures can give us an independent insight into perceived running speed. We found that the slope of the relationship between theta frequency and running speed was reduced within the VR compared to R, while this was not the case for the firing rates of place, grid and speed cells. See Figure 7. However, the changes in grid scale and the changes theta frequency in virtual versus real environments did not correlate with each other significantly across animals.

**Figure 7.**
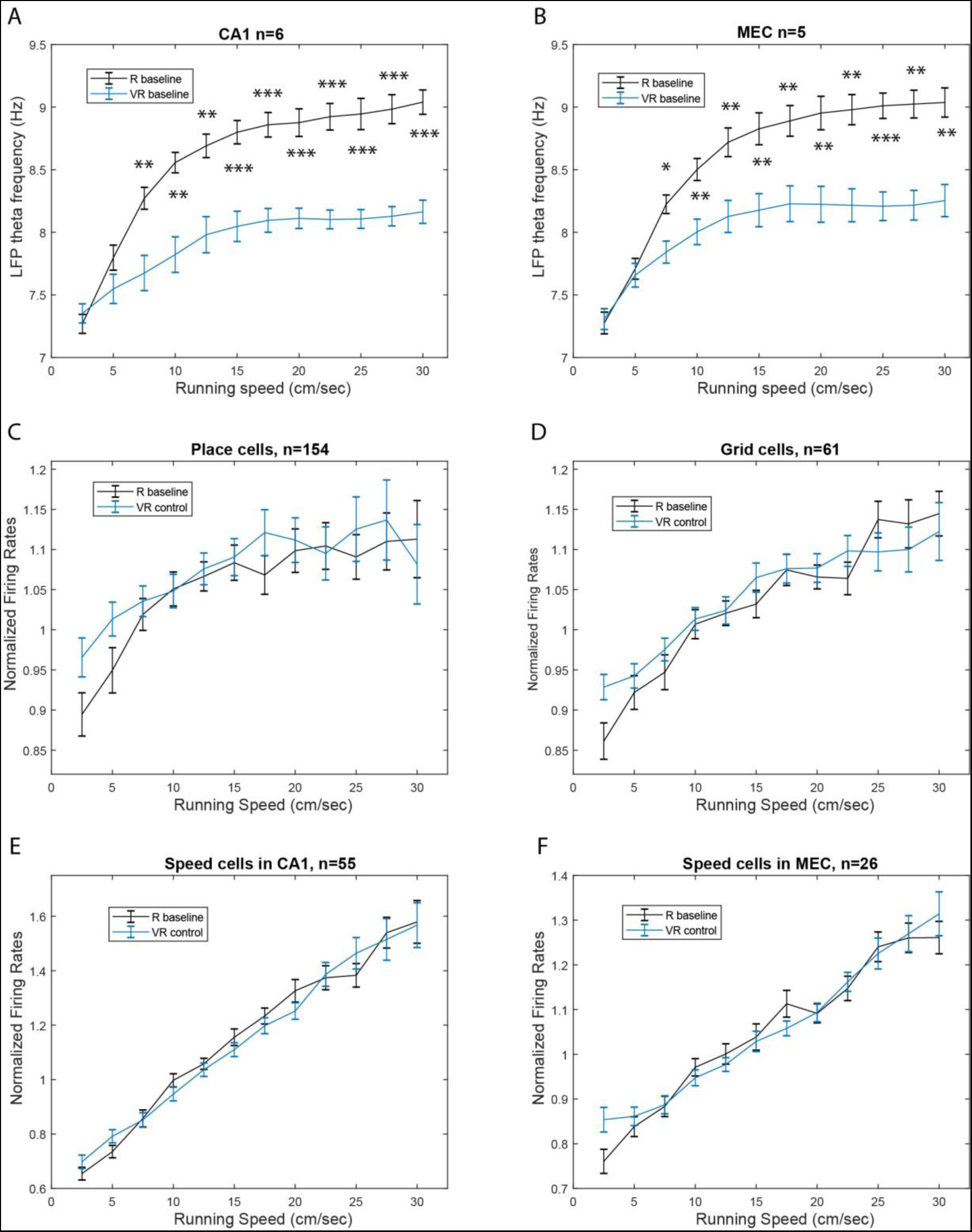
Effect of running speed on theta frequency and firing rates in real and virtual environments. Relationship between running speed in VR (blue) and R (black) on instantaneous LFP theta frequency in CA1 (A, n=6); instantaneous LFP theta frequency in dmEC (B, n=5); firing rates of place cells in CA1 (C, n=154); firing rates of grid cells in dmEC (D, n=61); speed-modulated cells in CA1 (E, n=55); firing rates of speed-modulated cells in dmEC (F, n=26). Lines show the mean (± s.e.m) theta frequency in each running speed bin (2.5cm/s to 30cm/s).

## Discussion

We have demonstrated the ability of a novel mouse virtual reality (VR) system to allow expression of spatial learning and of the characteristic spatially modulated firing patterns of place, grid and head-direction cells in open arenas. Thus it passes the first pre-requisite as a tool for studying the mechanisms behind the two dimensional firing patterns of these spatial cells, following previous systems for rats that also allow physical rotation of the animal^17,18^. Head-fixed or body-fixed VR systems have been very successful for investigating the one-dimensional spatial firing patterns of place cells^2-4,12,14-16^ or grid cells^6-9^, e.g. modulation of firing rate by the distance along a linear trajectory. But the two-dimensional firing patterns of place, grid or head direction cells are not seen in these systems.

Although the characteristic firing patterns of spatial cells were expressed within our VR system, there were also some potentially instructive differences in their more detailed properties between VR and a similar real environment (R), which we discuss below.

The spatial scale of the firing patterns of both place cells and grid cells was approximately 1.4 times larger in VR compared to R (Figures 3 and 4; see also^17^). Along with the increased scale of place and grid cell responses in VR, there was a reduction in the dependence of theta frequency on running speed. The LFP theta frequency reflects a contribution from vestibular translational acceleration signals^14,30^ which will be absent in our VR system. However, there was no change in the increasing firing rates of place, grid and speed cells with running speed in VR (Figure 7), indicating an independence from vestibular translational acceleration cues. Thus it is possible that the absence of linear acceleration signals affects both LFP theta rhythmicity and the spatial scale of firing patterns, but there was no evidence that the two were directly related.

Finally, uncontrolled distal cues, such as sounds and smells, and the visual appearance of the apparatus aside from the screens (the edge of the ball, the edges of the screens) will conflict with virtual cues indicating self-motion. Thus increased firing field size could also reflect broader tuning or reduced precision due to absent or conflicting inputs, consistent with the reduced spatial information seen in place and grid cell firing patterns (Figures 3 and 4), and potentially a response to spatial uncertainty^31^.

The head direction cells do not show broader tuning in the VR (Figure 5), probably because there is no absence of vestibular rotation cues and no conflict with distal real-world cues, as the mice rotate similarly in the virtual and real world. We note however, that spatial firing patterns follow the virtual cues when the virtual cues and entry point are put into conflict with uncontrolled real-world cues (Figure 6).

Place cell firing in VR showed an increased directionality compared to the real environment. One possible explanation, that the apparent directionality reflected inhomogeneous sampling of directions in the firing field, was not supported by further analyses (Figure 3E). A potential benefit of VR is an absence of local sensory cues to location, as experimenters typically work hard to remove consistent uncontrolled cues from real-world experiments (e.g., cleaning and rotating the walls and floor between trials). However, but reliable within-trial local cues may contribute to localisation of firing nonetheless^14^. Thus it maybe that uncontrolled local cues in real experiments (even if unreliable from trial to trial) are useful for supporting a locational response that can be bound to the distinct visual scenes observed in different directions, see also^15^. In this case, by removing these local cues, the use of VR leaves the locational responses of place cells more prone to modulation by the remaining (directionally specific) visual cues. We note that grid cells did not show such an increase in directional modulation as the place cells. This may indicate that place cell firing is more influenced by environmental sensory inputs - and thus directional visual inputs given the absence of local cues, while grid cell firing might be more influenced by self-motion cues, and thus less dependent on local cues for orientation independence. However, this would need to be verified in future work.

In conclusion, by using VR, the system presented here offers advantages over traditional paradigms by enabling manipulations that are impossible in the real-world, allowing visual projection of an environment that need not directly reflect a physical reality or the animals’ movements. Differences between the firing patterns in VR and R suggest broader spatial tuning as a possible response to under-estimated translation or spatial uncertainty caused by missing or conflicting inputs, a role for local cues in supporting directionally independent place cell firing and potentially for self-motion cues in supporting directionally independent grid cell firing. Finally, the differential effects of moving from R to VR on the dependence on running speed of the LFP theta frequency compared to neuronal firing rates suggests distinct mechanisms for speed coding, potentially reflecting a differential dependence on vestibular translational acceleration cues.

Previous body-rotation VR systems for rats^17,18^ also allow expression of the two dimensional firing patterns of place, grid and head-direction cells. However, by working for mice and by constraining the head to rotation in the horizontal plane, our system has the potential for future use with multiphoton imaging using genetically encoded calcium indicators. The use of multiple screens and floor projectors is not as elegant as the single projector systems^17,18^ but allows the possible future inclusion of a two photon microscope above the head without interrupting the visual projection, while the effects of in-plane rotation on acquired images should in principle be correctable in software.

## Materials and Methods

### Virtual Reality

A circular head-plate made of plastic (Stratasys Endur photopolymer) is chronically attached to the skull, with a central opening allowing the implant of tetrodes for electrophysiological recording (see Surgery). The head-plate makes a self-centring joint with a holder mounted in a bearing (Kaydon reali-slim bearing KA020XP0) and is clipped into place by a slider. The bearing is held over the centre of an air-supported Styrofoam ball. Four LCD screens placed vertically around the ball and two projectors onto a horizontal floor provide the projection of a virtual environment. The ball is prevented from yaw rotation to give the mouse traction to turn and to prevent any rotation of the ball about its vertical axis, following^17^. See Figure 1A-E.

The virtual environment runs on a Dell Precision T7500 workstation PC running Windows 7 64-bit on a Xeon X5647 2.93GHz CPU, displayed using a combination of four Acer B236HL LCD monitors mounted vertically in a square array plus two LCD projectors (native resolution 480×320, 150 lumens) mounted above to project floor texture. The head-holder is at the centre of the square and 60mm from the bottom edge of the screens, and 9500mm below the projectors. The LCD panels are 514mm × 293mm, plus bezels of 15mm all around. These six video feeds are fed by an Asus AMD Radeon 6900 graphics card and combined into a single virtual display of size 5760×2160px using AMD Radeon Eyefinity software. The VR is programmed using Unity3d v5.0.2f1 which allows virtual cameras to draw on specific regions of the virtual display, with projection matrices adjusted (see Kooima, 2008 http://csc.lsu.edu/~kooima/articles/genperspective/index.html) to the physical dimensions and distances of the screens and to offset the vanishing point from the centre. For example, a virtual camera facing the X-positive direction renders its output to a portion of the virtual display which is known to correspond to the screen area of the physical monitor facing the X-negative direction.

Translation in the virtual space is controlled by two optical mice (Logitech G700s gaming mouse) mounted with orthogonal orientations at the front and side of a 200mm diameter hollow polystyrene sphere, which floats under positive air pressure in a hemispherical well. The optical mice drive X and Y inputs respectively by dint of their offset orientations, and gain can be controlled within the Unity software. Gain is adjusted such that real-world rotations of the sphere are calibrated so that a desired environmental size (e.g. 600mm across) corresponds to the appropriate movement of the surface of the sphere under the mouse (i.e. moving 600mm, or just under one rotation, on the sphere takes the mouse across the environment). Mouse pointer acceleration is disabled at operating system level to ensure movement of the sphere is detected in a linear fashion independent of running speed.

The mouse is able to freely rotate in the horizontal plane, which has no effect on the VR display (but brings different screens into view). Rotation is detected and recorded for later analysis using an Axona dacqUSB tracker which records the position of two LEDs mounted at ~25mm offset to left and right of the head stage amplifier (see Surgery). Rotation is sampled at 50Hz by detection of the LED locations using an overhead video camera, while virtual location is sampled and logged at 50Hz.

Behaviour is motivated by the delivery of milk rewards (SMA, Wysoy) controlled by a Labjack U3HD USB Data Acquisition device. A digital-to-analogue channel applies 5V DC to a control circuit driving a 12V Cole-Parmer 1/16” solenoid pinch valve, which is opened for 100ms for each reward, allowing for the formation of a single drop of milk (5uL) under gravity feed at the end of a 1/32” bore tube held within licking distance of the animal’s mouth.

Control of the Labjack and of reward locations in the VR is via UDP network packets between the VR PC and a second experimenter PC, to which the Labjack is connected by USB. Software written in Python 2.7 using the Labjack, tk (graphics) and twistd (networking) libraries provide a plan-view graphical interface in which the location of the animal and reward cues in the VE can be easily monitored and reward locations manipulated with mouse clicks. See Figure 1.

### Animals

Subjects (11 C57Bl/6 mice) were aged 11-14 weeks and weighed 25-30 grams at the time of surgery. Mice were housed under 12:12 inverted light-dark cycle, with lights on at 10am. All work was carried out under the Animals (Scientific Procedures) Act 1986 and according to Home Office and institutional guidelines.

### Surgery

Throughout surgery, mice were anesthetized with 2-3% isoflurane in O2. Analgesia was provided preoperatively with 0.1mg/20g Carprofen, and post-operatively with 0.1mg/20g Metacam. Custom-made head plates were affixed to the skulls using dental cement (Kemdent Simplex Rapid). Mice were implanted with custom-made microdrives (Axona, UK), loaded with 17 μm platinum-iridium tetrodes, and providing buffer amplification. Two mice were implanted with 8 tetrodes in CA1 (ML:1.8mm, AP: 2.1mm posterior to bregma), three mice with 8 tetrodes in the dorsomedial entorhinal cortex (dmEC, ML = 3.1mm. AP = 0.2 mm anterior to the transverse sinus, angled 4° posteriorly), and six mice received a dual implant with one microdrive in right CA1 and one in left dmEC (each mircrodrive carried 4 tetrodes). After surgery, mice were placed in a heated chamber until fully recovered from the anaesthetic (normally about 1 h), and then returned to their home cages. Mice were given at least 1 week of post-operative recovery before cell screening and behavioural training started.

### Behavioural Training

Behavioural training in the virtual reality setup started while tetrodes were approaching target brain areas (see Screening for spatial cells). Behavioural training involved four phases. Firstly, mice experienced an infinitely long 10cm-wide virtual linear track, with 5 uL milk drops delivered as rewards. Reward locations were indicated by virtual beacons (high striped cylinders with a black circular base, see Figure S1A), which were evenly placed along the track (see Figure S1C). When the mouse contacted the area of the base, milk was released and the beacon disappeared (reappearing in another location). The lateral movement of the mice was not registered in this phase. The aim of this training phase was to habituate the mice to being head restrained and train them to run smoothly on the air-cushioned ball. It took three days, on average, for mice to achieve this criterion and move to the next training phase.

During the second training phase mice experienced a similar virtual linear track (see Figure S1B), which was wider than the first one (30cm wide). During this phase, reward beacons were evenly spaced along the long axis of the track, as before, but placed pseudo-randomly in one of three predefined positions on the lateral axis (middle, left or right). The aim of this training phase was to strengthen the association between rewards and virtual beacons, and to train animals to navigate towards rewarded locations via appropriate rotations on top of the ball. This training phase also took three days, on average. During the third training phase mice were introduced into a virtual square arena placed in the middle of a larger virtual room (see Figures 1E and S1C). The virtual arena had size 60×60cm or 90cm×90cm for different mice. Reward beacons had a base of diameter that equalled to 10% of the arena width. Mice were trained on a ‘random foraging’ task, during which visible beacons were placed in the square box at random locations (at any given time only one beacon was visible).

The last training phase was the ‘fading beacon’ task. During this task, every fourth beacon occurred in a fixed location (the three intervening beacons being randomly placed within the square enclosure; see Figure S1D). At the beginning of this training phase the ‘fixed location beacon’ slowly faded from view over 10 contacts with decreasing opacity. The beacon would remain invisible as long as mice could find it, but would became visible again if mice could not locate it after 2 min of active searching. Once mice showed consistent navigation towards the fading fixed beacon, they were moved to the ‘faded beacon’ phase of the task where the ‘fixed location beacon’ was invisible from the start of the trial and remained invisible throughout the trial, with two drops of milk given as reward for contact. This trial phase therefore requires mice to navigate to an unmarked virtual location starting from different starting points (random locations where the 3^rd^ visible beacon was placed). As such, the ‘fading beacon’ task serves like a continuous version of a Morris Water Maze task^19^, combining reference memory for an unmarked location with a foraging task designed to optimise environmental coverage for the assessment of spatial firing patterns. Mice typically experienced one trial per day.

### Behavioural analyses

All training trials in the VR square and the real (R, see Screening for spatial cells) square environments from the 11 mice were included in the behavioural analyses. During electrophysiological recording in R, the mouse’s position and head orientation were tracked by an overhead camera (50Hz sampling rate) using two infra-red LEDs attached to the micro-drive at a fixed angle and spacing (5 cm apart). Brief losses of LED data due to cable obstruction were corrected with linear interpolation between known position values. Interpolation was carried out for each LED separately. The position values for each LED were then smoothed, separately, using a 400ms boxcar filter. During electrophysiological recording in VR, head orientation was tracked as in R, the path, running speed and running direction was inferred from the VR log at 50Hz (movements of VR location being driven by the computer mice tracking the rotation of the ball, see above).

Path excess ratio was defined as the ratio between the length of the actual path that an animal takes to run from one reward location to another, and the distance between the two reward locations.

### Screening for spatial cells

Following recovery, mice were food restricted to 85% of their free-feeding body weight. They were then exposed to a recording arena every day (20 mins per day) and screening for neural activity took place. The recording arena was a 60×60cm square box placed on a black Trespa ‘Toplab’ surface (Trespa International B.V., Weert, Netherlands), and surrounded by a circular set of black curtains. A white cue-card (A0, 84 × 119 cm), illuminated by a 40 W lamp, was the only directionally polarising cue within the black curtains. Milk (SMA Wysoy) was delivered as drops on the floor from a syringe as rewards to encourage foraging behaviour. Tetrodes were lowered by 62.5 um each day, until grid or place cell activity was identified, in dmEC or CA1 respectively. Neural activity was recorded using DACQ (Axona Ltd., UK) while animals were foraging in the square environment. For further details see^2^.

### Recording spatial cell activity

Each recording session consisted of at least one 40-min random-foraging trial in a virtual square environment (see above for behavioural training). For 7 mice the virtual environment had size 60×60cm and for 4 mice 90×90cm when recording took place. After one (or more) 40-min random foraging trials in the virtual square, mice were placed in a real-world square (60×60cm square, similar to the screening environment, see above) for a 20-min random-foraging trial in real world.

Additionally, 4 mice also underwent a virtual cue rotation experiment, which consisted of two 40-min random-foraging VR trial (one baseline VR trial and one rotated VR trial) and one 20 min R trial. Two mice navigating 60×60 cm VR squares and two 90×90cm squares participated in this experiment. In the rotated VR trials, all cues in the virtual reality environment rotated 180 degrees compared to the baseline trial, as was the entry point mice were carried into the VR rig from.

### Firing rate map construction and spatial cell classification

Spike sorting was performed offline using an automated clustering algorithm (KlustaKwik^20^) followed by a manual review and editing step using an interactive graphical tool (waveform, Daniel Manson, http://d1manson.github.io/waveform/). After spike sorting, firing rate maps were constructed by binning animals’ positions into 1.5 × 1.5cm bins, assigning spikes to each bin, smoothing both position maps and spike maps separately using a 5×5 boxcar filter, and finally dividing the smoothed spike maps by the smoothed position maps.

Cells were classified as place cells if their spatial information in baseline trials exceeded the 99^th^ percentile of a 1000 shuffled distribution of spatial information scores calculated from rate maps where spike times were randomly offset relative to position by at least 4 sec. Cells were classified as grid cells if their gridness scores in baseline trials exceeded the 99^th^ percentile of a shuffled distribution of 1000 gridness scores^21^. Cells were classified as head direction cells if their Rayleigh vectors in baseline trials exceeded the threshold of the 99^th^ percentile population shuffling.

Speed-modulated cells were classified from the general population of the recorded cells following^22^. Briefly, the degree of speed modulation for each cell was characterised by first defining the instantaneous firing rate of the cell as the number of spikes occurring in each position bin divided by the sampling duration (0.02s). Then a linear correlation was computed between the running speeds and firing rates across all position samples in a trial, and the resulting r-value was taken to characterise the degree of speed modulation for the cell. To be defined as speed-modulated, the r-value for a cell had to exceed the 99th percentile of a distribution of 1000 r-values obtained from spike shuffled data.

When assessing the directional modulation of place and grid cell firing (Figures 3 and 4), apparent directional modulation can arise in binned firing rate data from heterogenous sampling of directions within the spatial firing field^23,24^. Accordingly we fit a joint (‘pxd’) model of combined place and directional modulation to the data (maximising the likelihood of the data^24^) and perform analyses on the directional model in addition to the binned firing rate data.

## Acknowledgements

We acknowledge support from the Wellcome Trust, European Union’s Horizon 2020 research and innovation programme (grant agreement No. 720270), Biotechnology and Biological Sciences Research Council, European Research Council and China Scholarship Council, and technical help from Peter Bryan, Duncan Farquharson and Daniel Manson.

## Supplementary Figures

Supplementary Video – example of a mouse performing the ‘fading beacon’ task.

**Figure S1.**
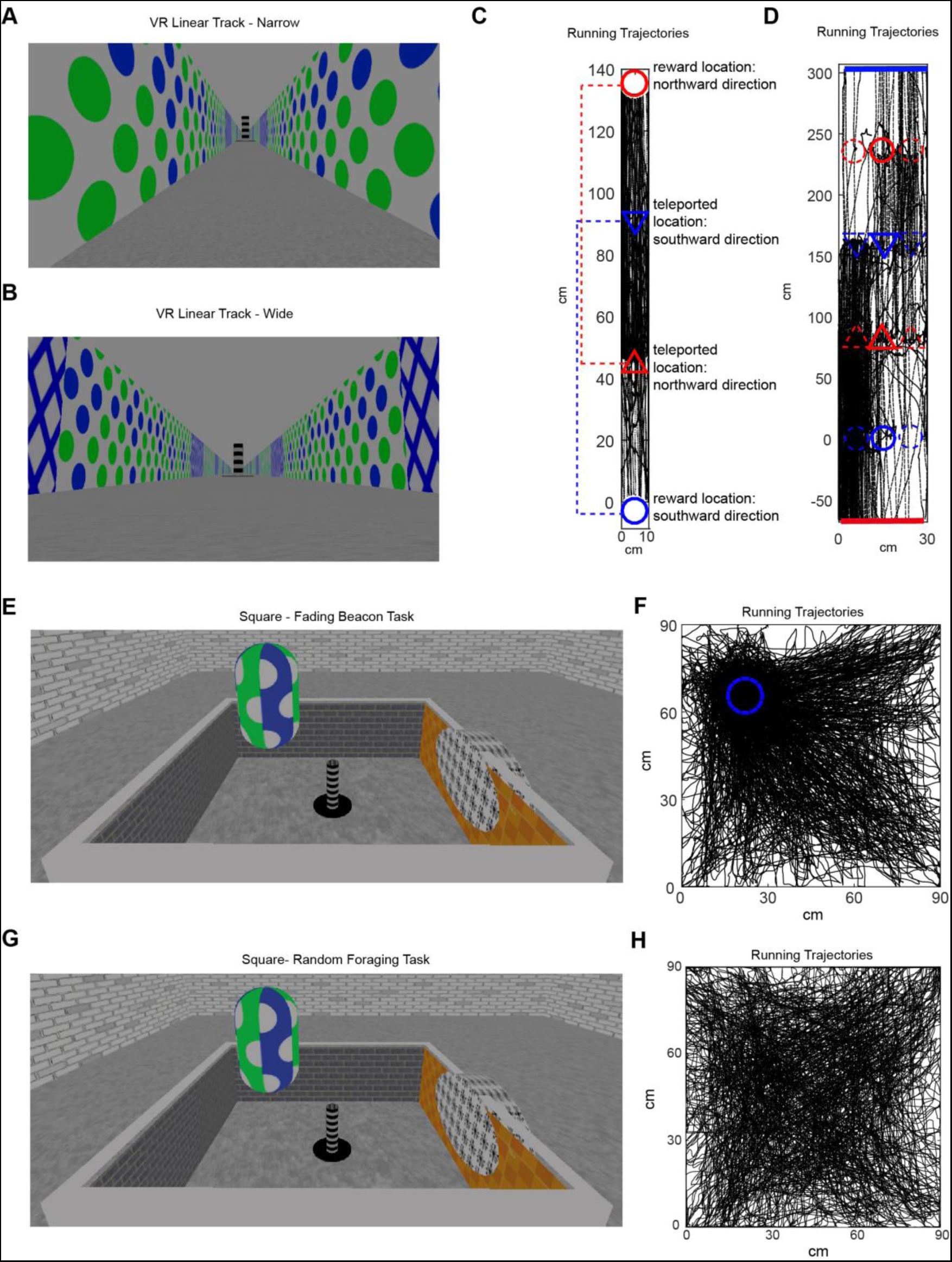
Example paths in the three training stages and the recording stage. (A) A view of the VR narrow linear track at training stage 1. (B) A view of the VR wide linear track at training stage 2. (C) An example running trajectory in the narrow linear track. Circles indicate reward locations, triangles indicate start points where animals get teleported after getting rewards. The start points were associated with the reward positions with matching colours (dotted lines). For example, a mouse gets teleported back to the position indicated by the blue triangle from the reward position indicated by the blue cycle. (D) An example running trajectory in the wide linear track. Circles indicate reward locations, bars indicate end points of the track, triangles indicate start points where animals get teleported after getting rewards or reaching the ends of the track. (E) A view of the virtual square at training stage 3 – the ‘fading beacon’ task. (F) An example trajectory when a mouse performing a ‘fading beacon’ task in the VR square. The dotted blue circle indicates the fixed location of every 4^th^ reward. (G) A view of the virtual square during the recording stage – random foraging. (H) An example trajectory from a mouse foraging for randomly-positioned rewards in the VR square.

**Figure S2.**
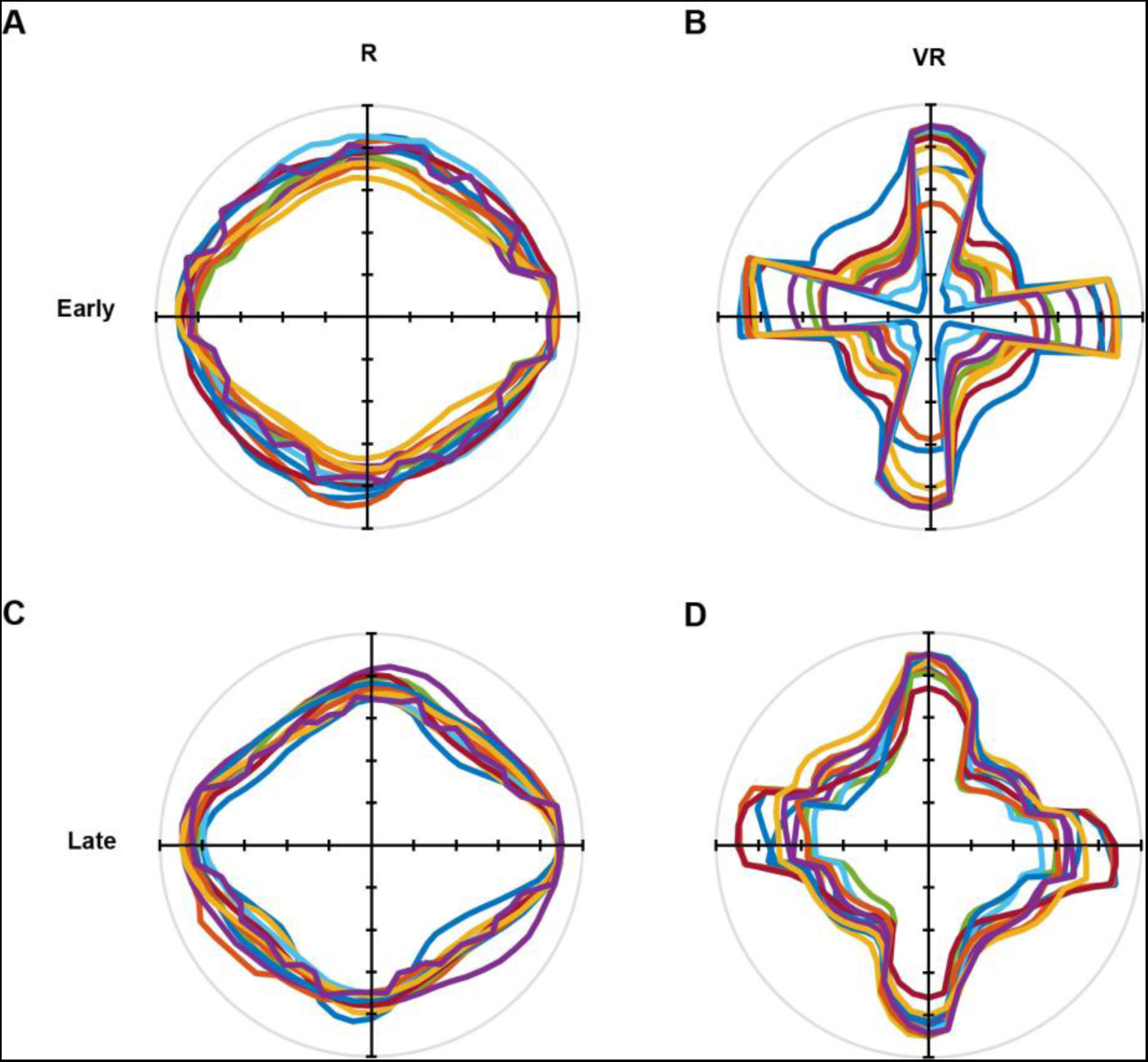
Directional polar plots of running directions (n=11 mice). (A-B) Average trajectories over the first 5 training trials in R (left column) and VR (right column). (C-D) Average trajectories over last 5 training trials in R (left column) and VR (right column). Each colour represents one animal.

**Figure S3.**
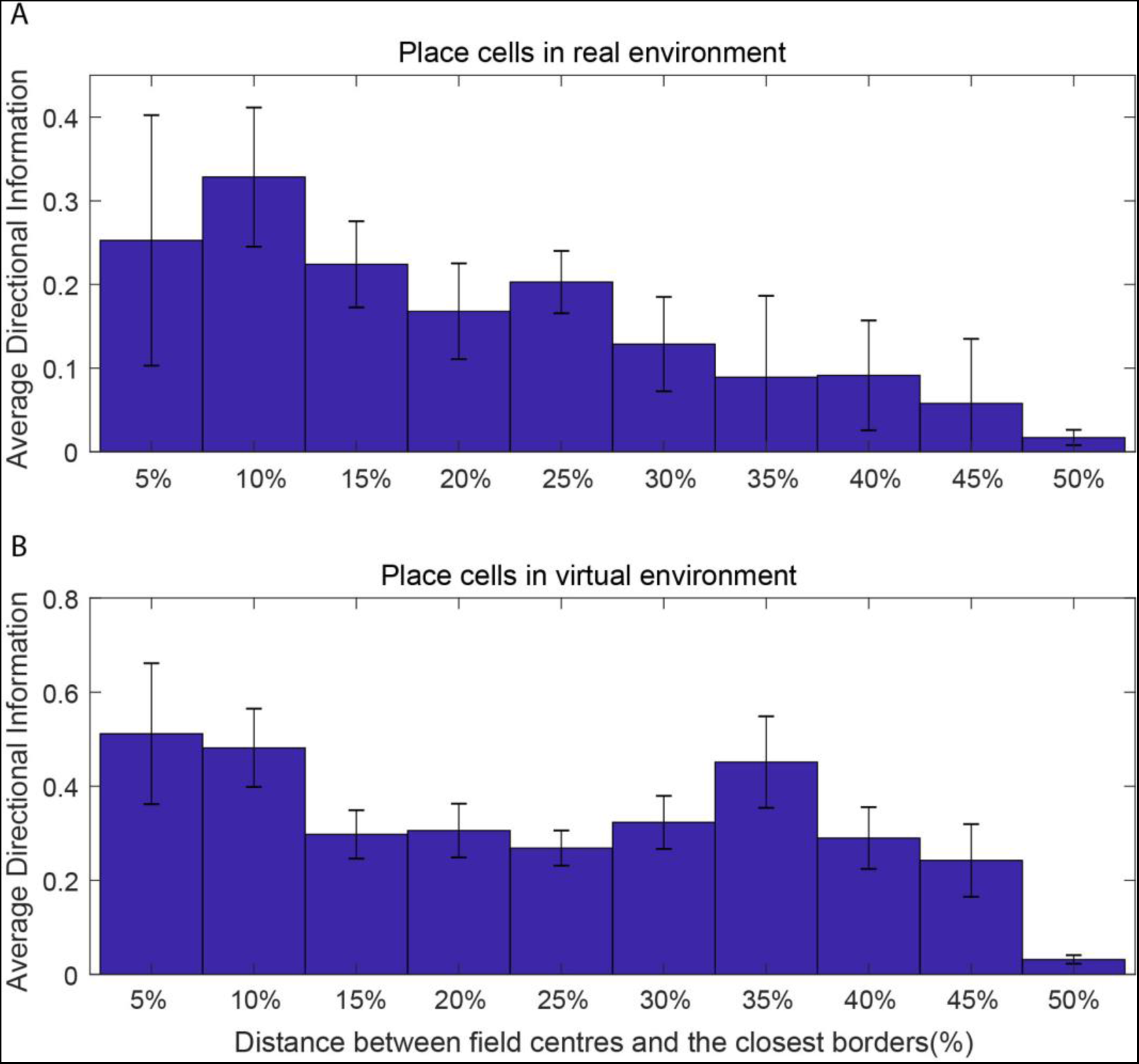
Directional information of place cell firing (bits/spike) as a function of the distance from the nearest wall (as % of the width of environment) in real (A) and virtual (B) environments.

**Figure S4.**
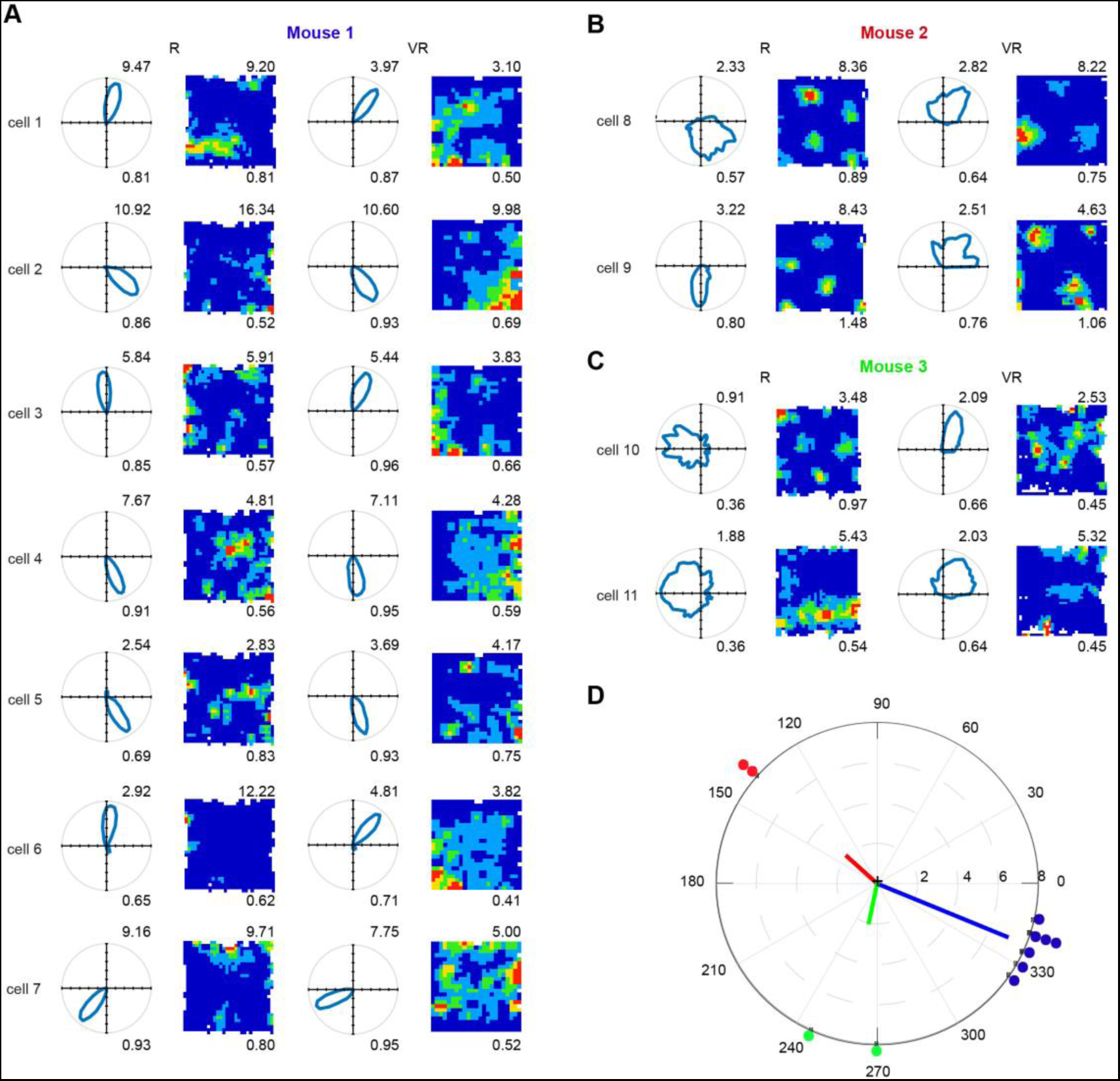
Eleven head direction cells recorded in dmEC. (A) Polar plots (left column) and firing rate maps (right column) of seven cells found in mouse 1. (B) Two conjunctive grid cells found in mouse 2. (C) Two cells found in mouse 3. Numbers on the top right show maximum firing rates, and on the bottom show Rayleigh vector length (left columns) and spatial information (right columns). (D) The relative directional tuning difference of simultaneously recorded head-direction cells between VR and R: Mouse 1 (blue), 337.71±8.28; Mouse 2 (red), 138.0±0.00; Mouse 3 (green), 258.00±16.97. The dots represent the relative directional tuning difference of individual cells between VR and R. The lines represent the mean tuning difference within the animals. Each dot represents one cell, and each colour represents one animal.

